# Alternative splicing of bicistronic *MOCS1* defines a novel mitochondrial protein maturation mechanism

**DOI:** 10.1101/429183

**Authors:** Simon Julius Mayr, Juliane Röper, Geunter Schwarz

## Abstract

Molybdenum cofactor biosynthesis is a conserved multistep pathway. The first step, the conversion of GTP to cyclic pyranopterin monophosphate (cPMP), requires bicsistronic *MOCS1*. Alternative splicing of *MOCS1* in exons 1 and 9 produces four different N-terminal and three different C-terminal products (type I-III). Type I splicing results in bicistronic transcripts with two open reading frames, of which only the first, MOCS1A, is translated, whereas type II/III splicing produces two-domain MOCS1AB proteins. Here, we report and characterize the mitochondrial translocation of alternatively spliced MOCS1 proteins. While MOCS1A requires exon 1a for mitochondrial translocation, MOCS1AB variants target to mitochondria via an internal motif overriding the N-terminal targeting signal. Within mitochondria, MOCS1AB undergoes proteolytic cleavage resulting in mitochondrial matrix localization of the MOCS1B domain. In conclusion we found that *MOCS1* produces two functional proteins, MOCS1A and MOCS1B, which follow different translocation routes before mitochondrial matrix import, where both proteins collectively catalyze cPMP biosynthesis. MOCS1 protein maturation provides a novel mechanism of alternative splicing ensuring the coordinated targeting of two functionally related mitochondrial proteins encoded by a single gene.

## Introduction

The molybdenum cofactor (Moco) is found in all kingdoms of life forming the active center of molybdenum-containing enzymes, except nitrogenase. In mammals, there are four enzymes that depend on Moco: sulfite oxidase, xanthine oxidase, aldehyde oxidase and the mitochondrial amidoxime-reducing component (Schwarz et al., 2009). Moco is composed of an organic pterin moiety that binds molybdenum via a dithiolene group attached to a pyran ring, which is synthesized by a complex biosynthetic pathway. A mutational block in any step of Moco biosynthesis results in Moco deficiency (MoCD), a severe inborn error of metabolism characterized by rapidly progressing encephalopathy and early childhood death (Schwarz, 2005). In recent years, efficient treatment for Moco-deficient patients with a defect in the first step of Moco-biosynthesis has been established (Veldman et al., 2010, Schwahn et al., 2015).

Moco biosynthesis is divided into three major steps, starting with GTP followed by the synthesis of three intermediates: cyclic pyranopterin monophosphate (cPMP) (Santamaria-Araujo et al., 2004), molybdopterin or metal-binding pterin (MPT) (Schwarz, 2005) and adenylated MPT (Kuper et al., 2004). Due to the strict evolutionary conservation of the pathway, intermediates are identical in all kingdoms of life. The first and most complex reaction sequence is catalyzed by two proteins in bacteria (MoaA and MoaC) while in humans different translation products originating from different open reading frames (ORFs) of the *MOCS1* gene (MOCS1A and MOCS1AB) are required for cPMP synthesis (Stallmeyer et al., 1999, Wuebbens et al., 2000, Hanzelmann et al., 2002).

*E. coli* MoaA and mammalian MOCS1A belong to the superfamily of radical S-adenosylmethionine (SAM) proteins with two highly oxygen-sensitive [4Fe-4S] clusters. The respective reaction mechanism has been studied for MoaA and involves reductive cleavage of SAM by the N-terminal [4Fe-4S] cluster (Hanzelmann et al., 2004). The resulting 5’-deoxyadenosyl radical initiates the transformation of 5’-GTP, which is bound to a C-terminal [4Fe-4S] cluster, by abstracting the 3’ proton from the ribose. Following a multi-step rearrangement reaction, 3,8’cH_2_-GTP is released (Hover & Yokoyama, 2015) and further processed by the second protein (*E. coli* MoaC, respectively human MOCS1AB) (Hanzelmann et al., 2002) resulting in pyrophosphate release and cyclic phosphate formation (Fig. 1A) (Wuebbens et al., 2000, Hover et al., 2015).

**Figure 1.**
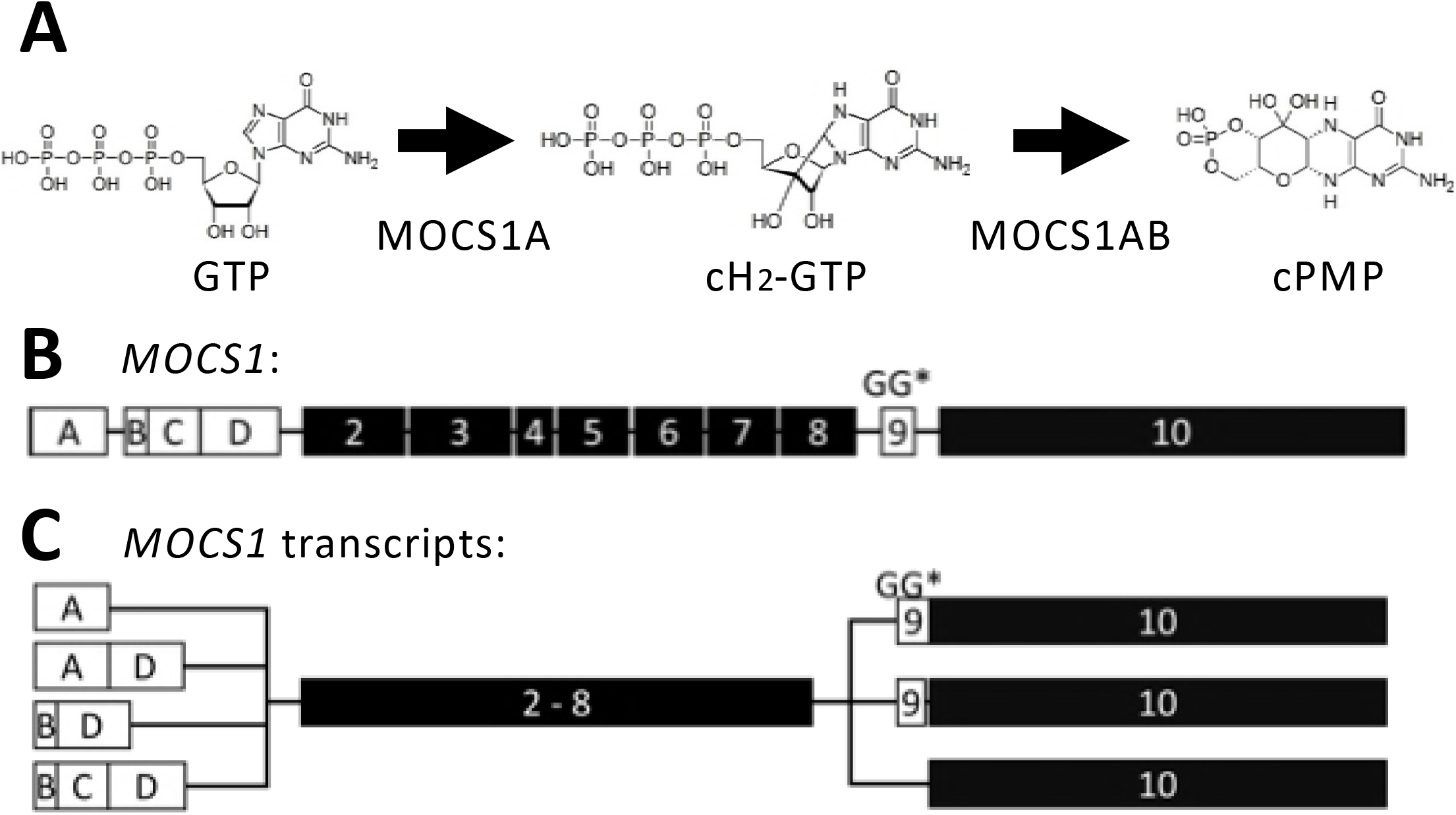
Alternative splicing of the *MOCS1* gene and gene products involved in cPMP synthesis. A) Biosynthesis of cPMP by the MOCS1 proteins in a two sub-step reaction. The MOCS1A protein converts GTP to the cyclic-dihydro-GTP intermediate, followed by the conversion of the intermediate to cPMP by MOCS1AB. B) Exon structure of the *MOCS1* gene highlighting the four alternatively splice exon 1 cassettes (white boxes), the shared exons 2–8 (Black boxes), the alternatively spliced exon 9 (white box) encoding the C-terminal double glycine motif (GG) and stop codon (*), as well as the MOCS1B encoding exon 10 (black box). Introns are indicated by a black line but are not reflecting actual intron size. C) Alternative splicing of *MOCS1* resulting in 12 different MOCS1 proteins. The splicing of exon 1 creates four alternative combinations (white boxes), while exon 9 splicing produces either the bicistronic type I variants when exon 9 (white box) is not spliced out leaving the double glycine motif and the stop codon in place. As a consequence active MOCS1A (exons 1–9), but not MOCS1B (exon 10) is translated. The monocistronic type II and III variants on do not contain the double glycine motif and the stop codon by having exon 9 removed either partially (type II) or entirely (type III), resulting in a MOCS1AB fusion protein representing active MOCS1B.

Interestingly, in the first two steps of Moco biosynthesis (cPMP and MPT synthesis) bicistronic transcripts (*MOCS1* and *MOCS2*) have been reported (Reiss et al., 1998a, Stallmeyer et al., 1999), which is very unusual in human gene expression (Lu et al., 2013). Proteins involved in cPMP synthesis are encoded by the *MOCS1* gene harboring 10 exons (Fig. 1B) leading to different alternatively spliced *MOCS1* transcripts (Gray & Nicholls, 2000), which are classified into three forms: Type I transcripts are bicistronic mRNAs with two non-overlapping ORFs, MOCS1A and MOCS1B (Reiss et al., 1998b), of which only the first ORF is translated yielding active MOCS1A. Type II and III transcripts are derived from two alternative splice sites within exon 9, both resulting in the lack of the MOCS1A stop codon and an in-frame-fusion with the second ORF (MOCS1B) thus producing a monocistronic transcript. Consequently, MOCS1B is not expressed independently. Splice type II only lacks 15 nucleotides of exon 9, whereas in the type III variant the entire exon 9 is absent (Fig. 1C). The resulting MOCS1AB fusion proteins harbor an active MOCS1B domain and a catalytically inactive MOCS1A domain due to deletion of the last 2 residues harboring a conserved C-terminal double-glycine motif (Hanzelmann et al., 2002). Type I and type III variants present 41 % and 55 % of all MOCS1A transcripts, respectively, the remainder being type II transcripts (Arenas et al., 2009).

In addition to exon 9 alternative splicing, cDNA sequences with four different 5’-regions have been reported and were named after the first authors describing the respective variants. Two alternative ATG start codons are encoded either by exon 1a (present in LARIN and ARENAS variants) or exon 1b (present in REISS and GROSS variants). Due to the lack of an intronic 3’ splice site in LARIN, exon 1a is spliced to exon 1d, while in ARENAS it is directly joined to exon 2 (subsequently referred to as MOCS1-ad and MOCS1-a). In a third REISS variant, exon 1b is joined with exons 1c and 1d, while in GROSS, it is spliced together with exon 1d (MOCS1-bcd and MOCS1-bd)(Fig. 1C). MOCS1-ad and MOCS1-bcd transcripts were found in different organs, while expression levels of MOCS1-bd were found to be very low (Gross-Hardt & Reiss, 2002).

MoCD-causing mutations have been found in *MOCS1, MOCS2, MOCS3* and *GPHN* genes (Reiss et al., 2001, Reiss & Hahnewald, 2011, Huijmans et al., 2017). Compared to their bacterial orthologues, human MOCS1 proteins exhibit N-terminal extensions of up to 56 residues depending on the length of the N-terminal splicing product of *MOCS1* exon 1. In contrast, in plants two genes (*Cnx2* and *Cnx3*) encode for proteins required for cPMP synthesis, both of which are characterized by extension of up to 112 residues encoding for classical N-terminal mitochondrial targeting signals (Teschner et al., 2010). Therefore, we proposed that MOCS1 proteins also localize to mitochondria and asked the question how alternative splicing of *MOCS1* transcripts controls the function and localization of MOCS1 proteins.

We found that in the bicistronic *MOCS1* transcripts (MOCS1A) exon 1 splicing results in translocation to the mitochondrial matrix when exon 1a is translated, while exon 1b variants remain cytosolic. In contrast, all monocistronic transcripts (MOCS1AB) produced proteins that were imported into mitochondria, regardless of their exon 1 composition. Additional submitochondrial localization studies of the MOCS1AB proteins revealed that only the proteolytic MOCS1B cleavage product was imported into the mitochondrial matrix, while full-length MOCS1AB could only be witnessed on the outer mitochondrial membrane.

## Results

### Localization of MOCS1A proteins

Given the N-terminal extension of the MOCS1 proteins compared to the homologous bacterial MoaA protein, first the different N-terminal splice variants were investigated concerning their cellular localization, knowing that N-terminal extensions may be involved cellular translocation processes. Based on the published sequences for MOCS1-a (Arenas et al., 2009), MOCS1-ad (AF034374)(Reiss & Hahnewald, 2011), MOCS1-bd (Gross-Hardt & Reiss, 2002) and MOCS1-bcd (Reiss et al., 1998a) variants, the four exon 1 type I splice variants were created by fusion PCR and inserted into pEGFP-N1 and expressed in COS7 cells as EGFP fusion proteins. While the MOCS1A-ad-EGFP protein colocalized with the mitochondrial marker (Mitotracker)(Fig. 2A), a diffuse cytosolic distribution was observed for MOCS1A-bcd-EGFP (Fig. 2B), suggesting that exon 1a facilitates mitochondrial import, given that exon 1d and exons 2–9 are shared between MOCS1A-ad and MOCS1A-bcd. In accordance no difference in cellular localization between MOCS1A-a and MOCS1A-ad, as well as MOCS1A-bd and MOCS1A-bcd, could be observed (Fig. EV1).

**Figure 2.**
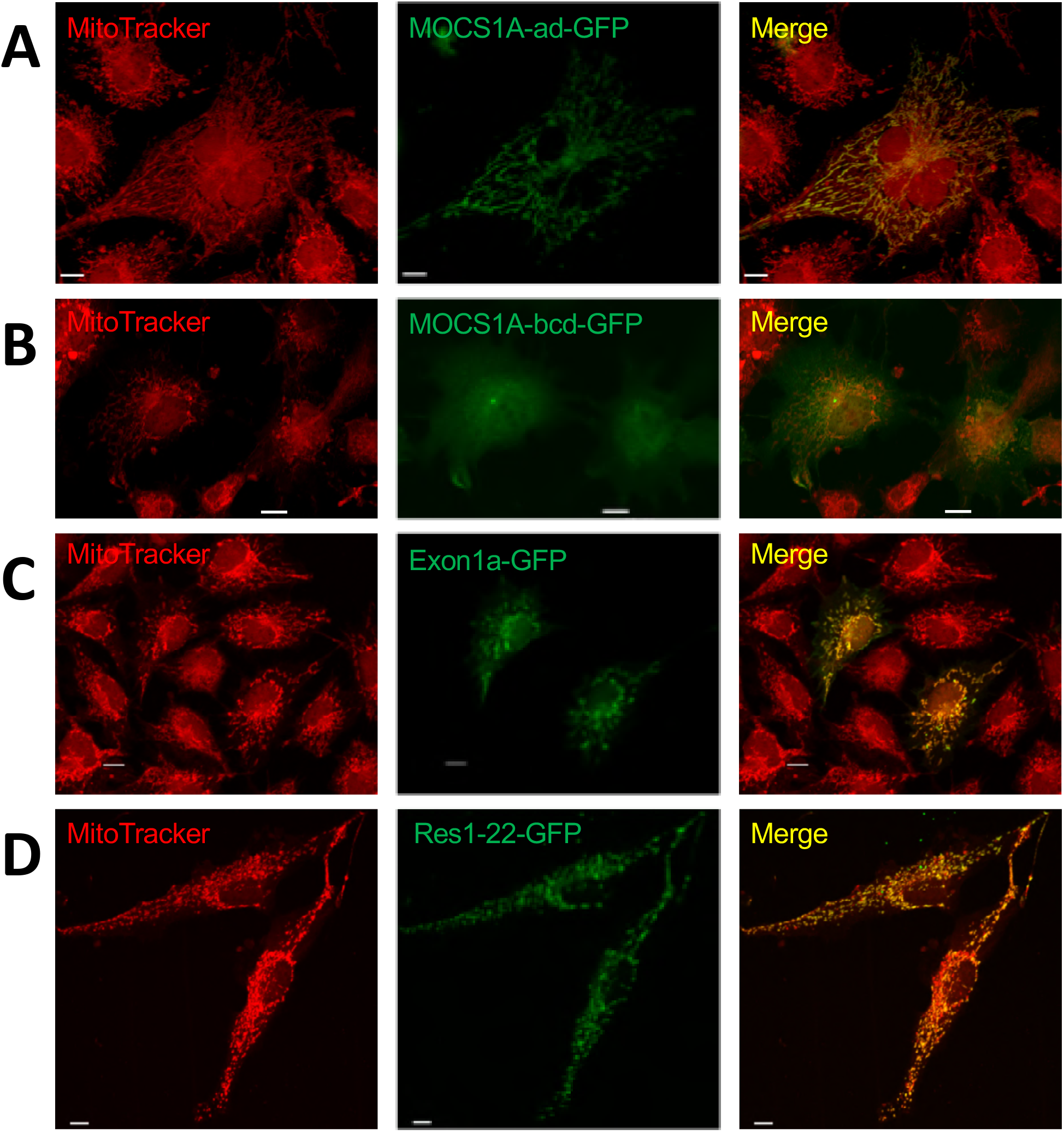
Transient expression of MOCS1A splice variants and MOCS1 derived constructs as GFP fusions in COS7 cells. COS7 cells were transfected with A) MOCS1A-ad, B) MOCS1A-bcd, C) exon 1a and D) residues 1–22 encoded by exon 1a. Following 48 h of transient expression, mitochondria were stained with MitoTracker®Red CMXRos and analyzed by confocal laser scanning microscopy. Overlay between the red and green (GFP) channel is shown in the yellow merge panel. Scale bars: 10 µm

**Extended View figure 1.**
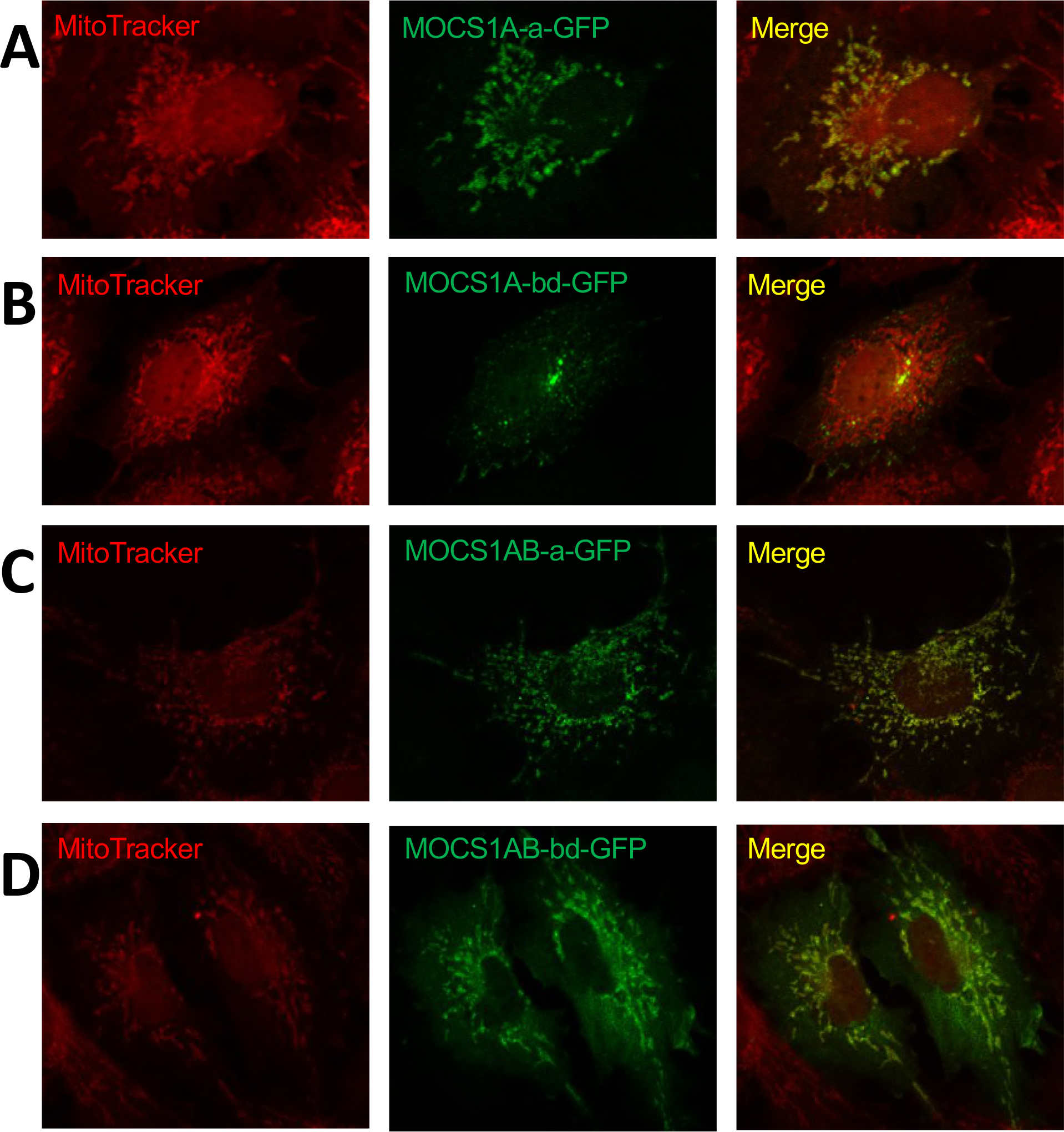
Transient expression of MOCS1A and MOCS1AB (type III) splice variants as GFP fusions in COS7 cells. COS7 cells were transfected with MOCS1A-a (A), MOCS1A-bd (B), MOCS1AB-a (C) and MOCS1AB-bd (D). Following 48 h of transient expression, mitochondria were stained with MitoTracker®Red CMXRos and analyzed by confocal laser scanning microscopy. Overlay between the red and green (GFP) channel is shown in the yellow merge panel.

To probe the role of exon 1a in mitochondrial targeting of MOCS1A, exon 1a was expressed as an EGFP fusion protein in COS7 cells resulting again in colocalization with the mitochondrial marker (Fig. 2C). Subsequent *in silico* analysis of the MOCS1A-ad sequence for mitochondrial translocation signals (MitoProtII)(Claros & Vincens, 1996) revealed a high mitochondrial import probability (99.94%) and a cleavage site after 22 residues (Fig. 3A). Subsequent expression of a fusion protein consisting of the N-terminal 22 residues and EGFP indeed resulted in colocalization of EGFP and the mitochondrial marker (Fig. 2D), confirming the presence of a classical N-terminal import signal in exon 1a. *In silico* analysis of the MOCS1A-ad sequence using heliquest (Gautier et al., 2008) revealed the formation of an amphipathic helix within the N-terminal 22 residue between Arg4 and Cys21 (Fig. 3A).

**Figure 3.**
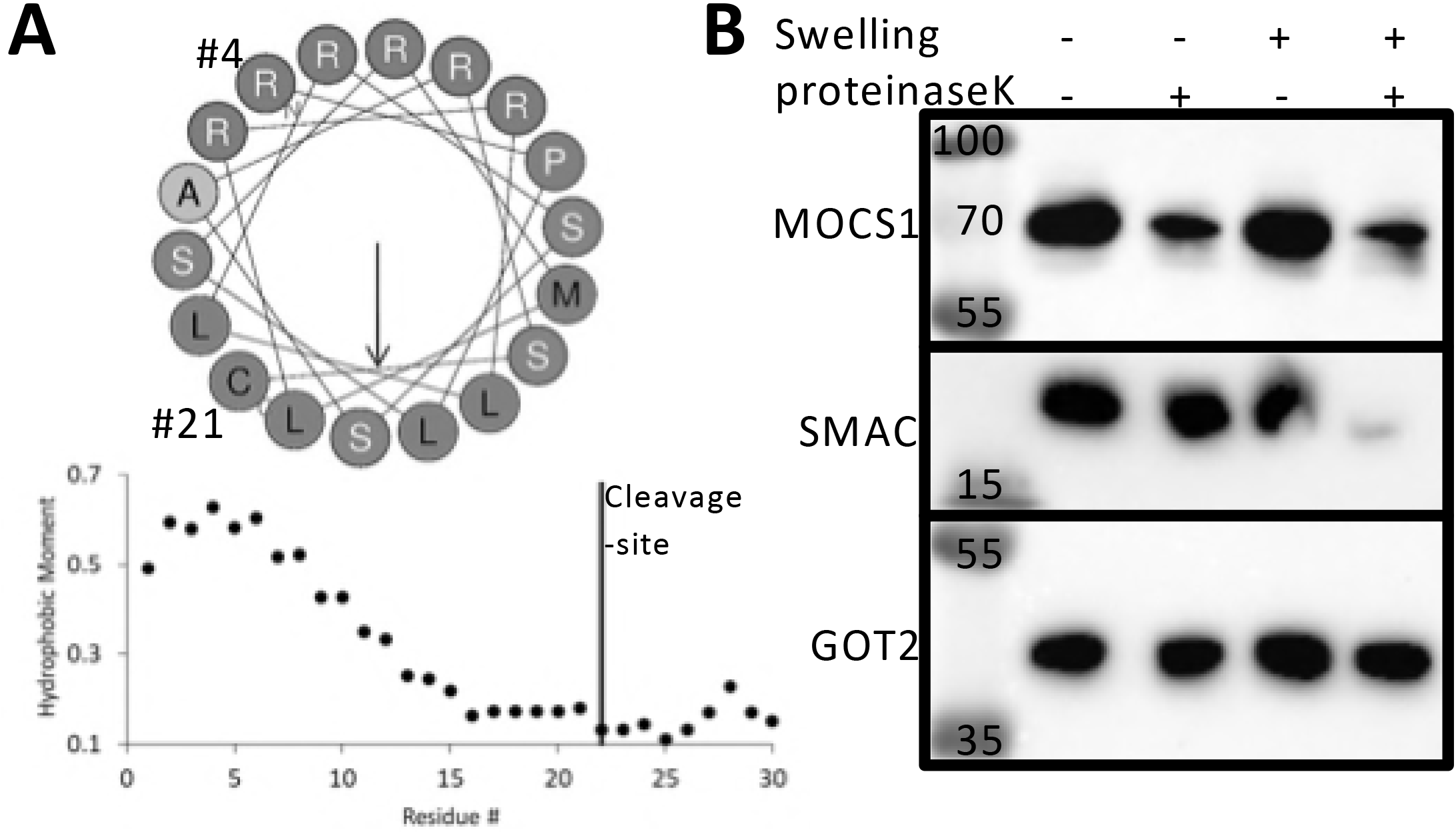
Mitochondrial import of MOCS1A. A) Prediction of N-terminal amphipathic helix consisting of the residues 4 to 21 encoded by exon 1a (heliquest). B) Partial proteolysis experiment of mitochondria enriched from HEK293 cells overexpressing MOCS1A-ad as a GFP fusion protein. One of each fraction of intact and swollen mitochondria were treated with proteinase K. Subsequent 10%-SDS-PAGE followed by western blot analysis was performed using anti-MOCS1, anti-SMAC and anti-GOT2 antibodies.

MOCS1A proteins harbor two [4Fe4S], of which the N-terminal cluster is coordinated by two cysteines encoded by exon 1d. Expression of all four MOCS1A isoforms yielded in comparable activities of all splice variants except MOCS1A-a, which showed strongly reduced activity as expected due to the absence of exon 1d (Fig. EV2). This finding underlines the importance of fully coordinated clusters pointing towards an import into the mitochondrial matrix representing a possible compartment of iron-sulfer cluster proteins, besides the cytosol and the nucleus (Ciofi-Baffoni et al., 2018). In order to probe for matrix import of MOCS1A-ad, we expressed the MOCS1A-ad-EGFP fusion protein in HEK293 cells and enriched mitochondria after two days of culture. Enriched mitochondria were resuspended in two different buffers either keeping the mitochondria stable or inducing hypotonic swelling and thereby disrupting the outer mitochondrial membrane. Addition of proteinase K to one aliquot of each condition revealed that proteinase K was not able to digest MOCS1A-ad in both intact and swollen mitochondria (Fig. 3B), indicating mitochondrial matrix localization, as confirmed by the efficient digestion of SMAC protein (IMS loading control) in the proteinase K treated swollen mitochondria sample, but not of the GOT2 protein (matrix loading control). As a result, MOCS1A was found to be translocated to the mitochondrial matrix via a classical mitochondrial targeting signal located in exon 1a.

**Extended View figure 2.**
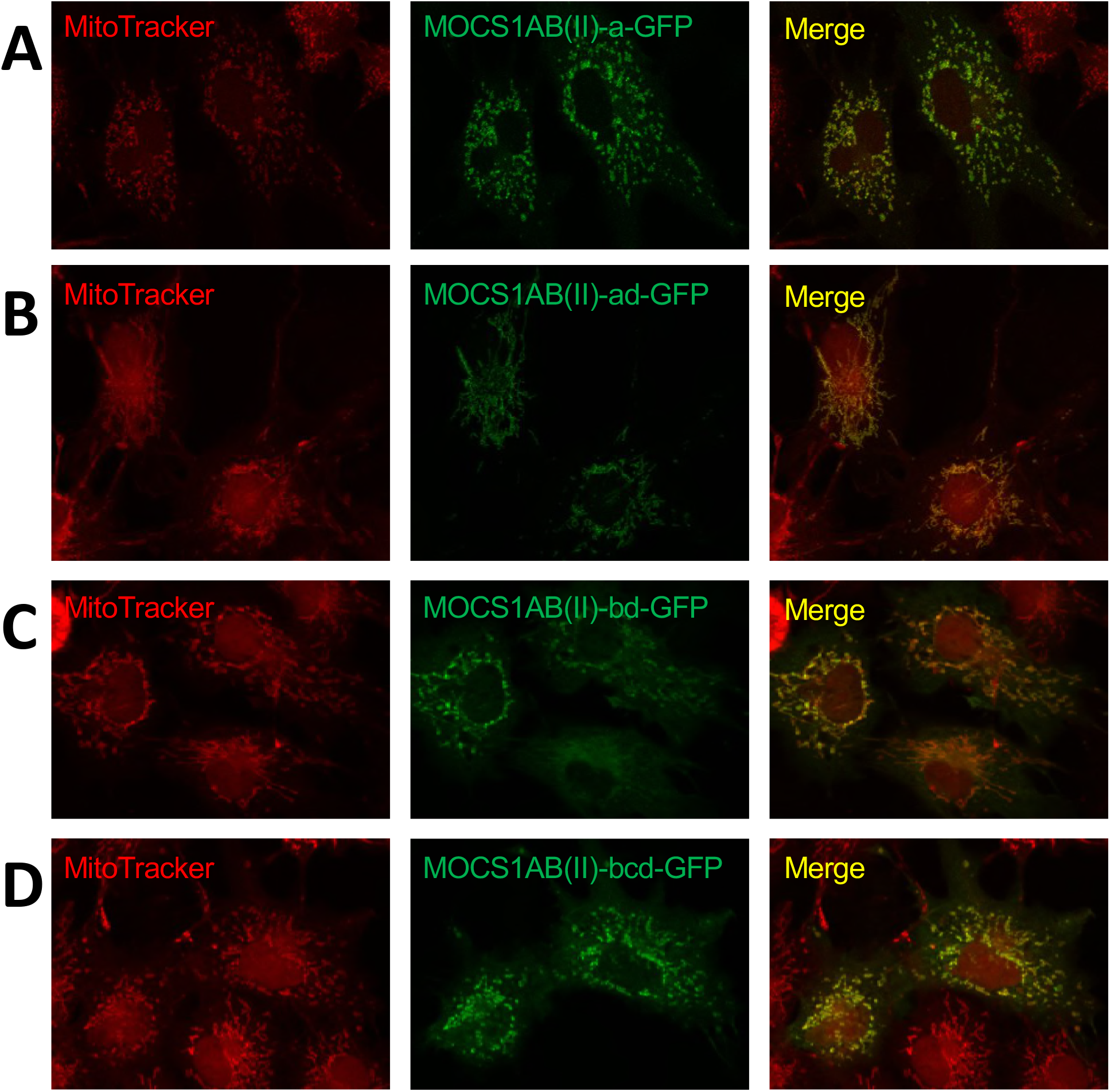
Functional reconstitution of Moco-deficient E. coli mutants KB 5242 (moaA–)with the respective MOCS1 proteins. HPLC analysis of Compound Z (cPMP oxidation product) in crude cell extracts. Measurements were performed in triplicate and standard deviations of at least three independent experiments with triplicate measurements are indicated by the error bars. Statistical analysis using the t-test: *** p ≤0.001, ** p≤ 0.01 and *p≤ 0.05.

### Localization of MOCS1AB proteins

Following the results obtained for the MOCS1A proteins, we next investigated the cellular localization of MOCS1AB proteins (splice type III, encoded by exons 1–8 and exon 10). Similar to the experimental design for MOCS1A proteins, we expressed MOCS1AB-ad-EGFP and MOCS1AB-bcd-EGFP in COS7 cells. As expected, MOCS1AB-ad-EGFP colocalized with the mitochondrial marker (Fig. 4A), however, when we expressed the MOCS1AB-bcd-EGFP fusion protein we again observed mitochondrial localization (Fig. 4B), even though exon 1a encoded residues were not present. Testing splice type II variants resulted in mitochondrial localization independent of exon 1 composition as well (Fig. EV3). Since the exons 1–8 are shared between the type I (MOCS1A) and type III (MOCS1AB) splice variants our finding suggested the presence of an additional translocation signal encoded by exon 10.

**Figure 4.**
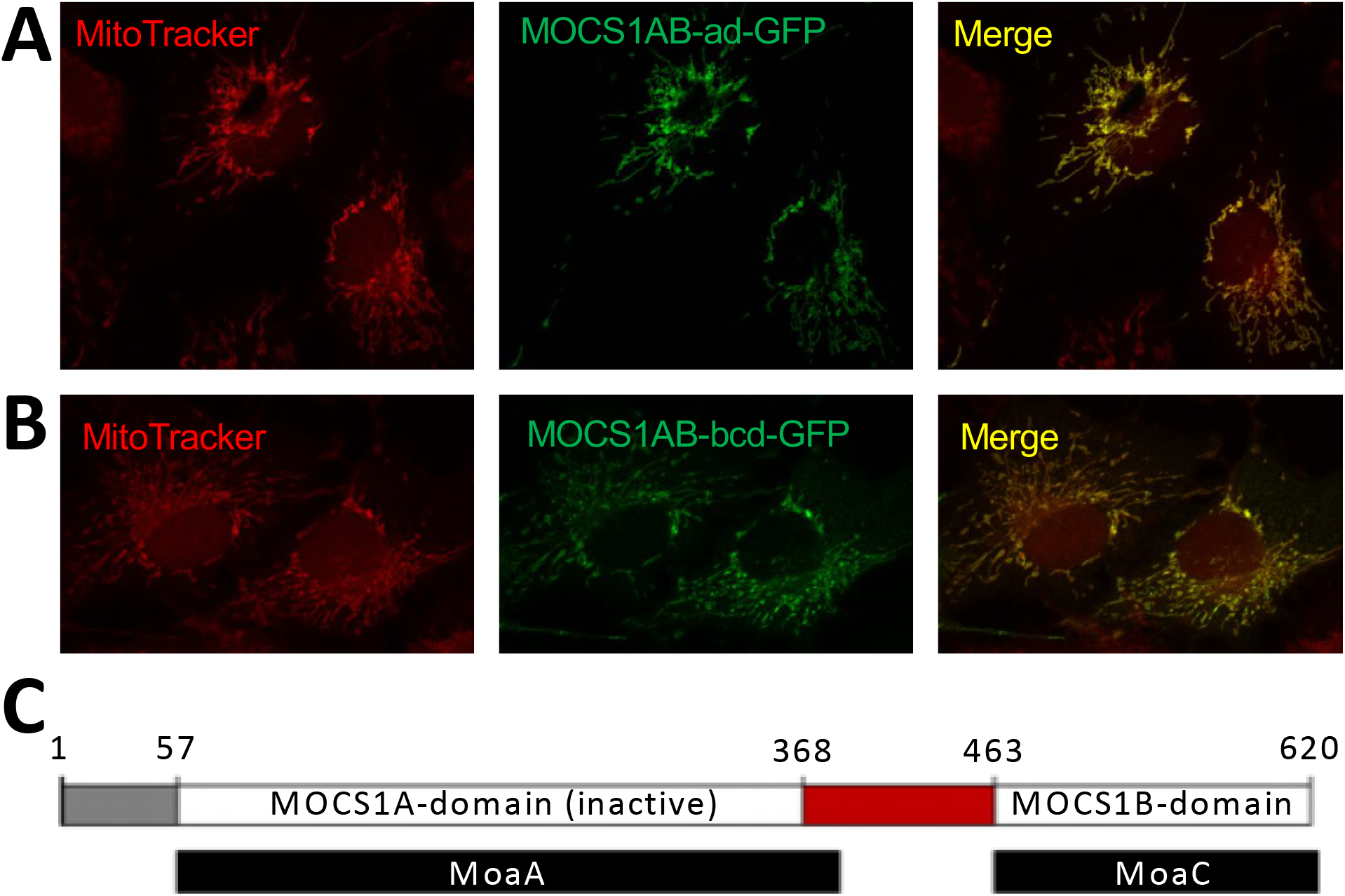
Identification of internal translocation signal of MOCS1AB proteins. A,B) Transient expression of MOCS1AB splice variants as GFP fusions in COS7 cells. COS7 cells were transfected with A) MOCS1AB-ad and B) MOCS1A-bcd. Following 48 h of transient expression, mitochondria were stained with MitoTracker®Red CMXRos and analyzed by confocal laser scanning microscopy. Overlay between the red and green (GFP) channel is shown in the yellow merge panel. C) Alignment of MOCS1AB (white box) with *E. coli* MoaA and MoaC (black boxes). Numbers indicate the positions of residues defining the N-terminal extension (gray box) and the internal extension (red box).

**Extended View figure 3.**
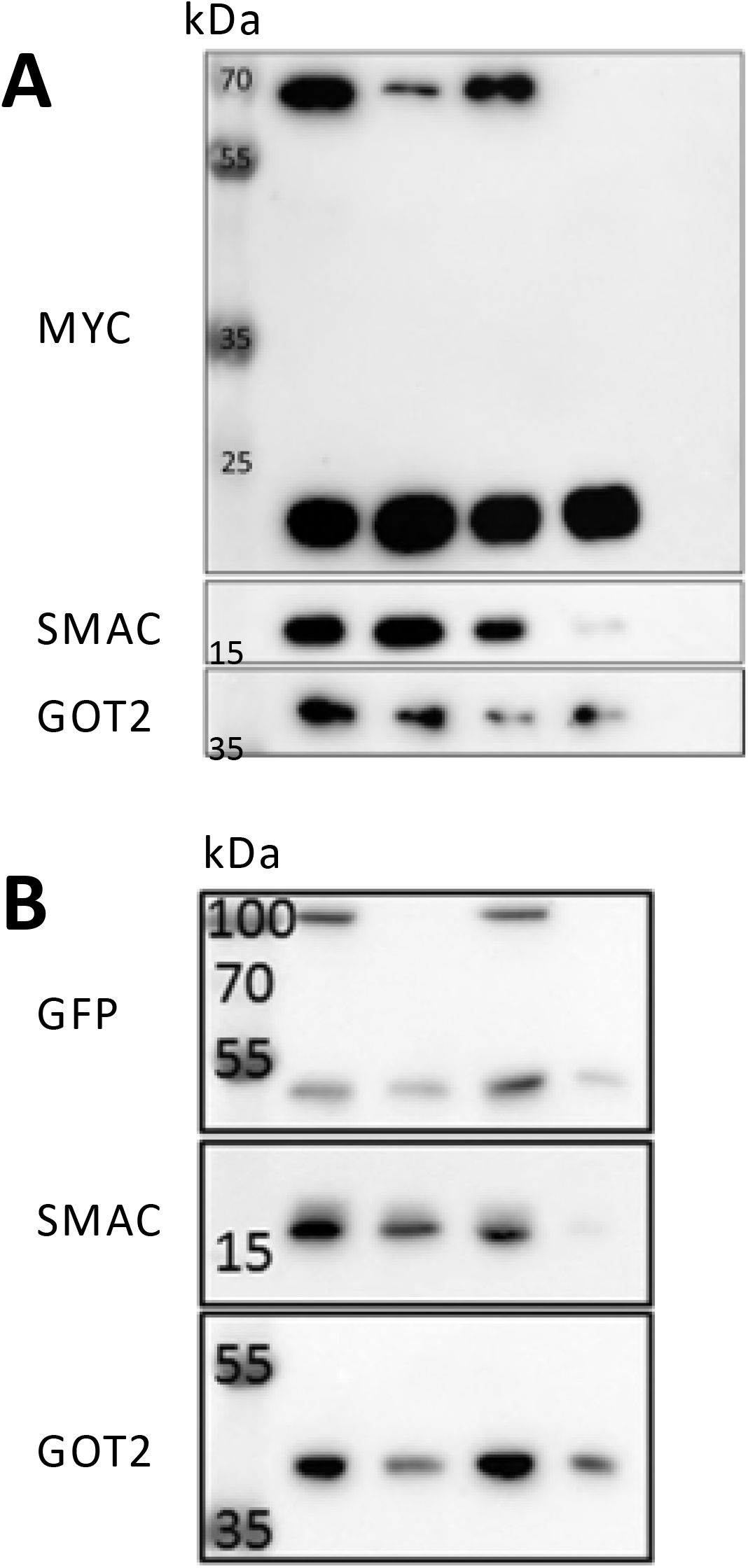
Transient expression of MOCS1AB (type II) splice variants as GFP fusions in COS7 cells. COS7 cells were transfected with MOCS1AB-a (A), MOCS1AB-ad (B), MOCS1AB-bd (C) and MOCS1AB-bcd (D). Following 48 h of transient expression, mitochondria were stained with MitoTracker®Red CMXRos and analyzed by confocal laser scanning microscopy. Overlay between the red and green (GFP) channel is shown in the yellow merge panel.

Alignment of MOCS1AB with the homologous bacterial proteins MoaA and MoaC revealed a polypeptide sequence of 95 non-conserved residues located between the MOCS1A and MOCS1B domain, encoded by the 5’ sequence of exon 10 (Fig. 4C). Such a sequence outside the functional domain might indicate an internal translocation signal, which was indeed confirmed by a stepwise TargetP *in silico* sequence analysis (Emanuelsson et al., 2000, Backes et al., 2018). Furthermore, subsequent heliquest sequence analysis showed the presence of another amphipathic helix resulting in a hydrophobic moment at position 395 (Gautier et al., 2008)(Fig. 5A). To probe this bioinformatic approach, full-length exon 10 (representing MOCS1B) and a deletion of the 5’ 285 bases of exon 10 (representing MOCS1B∆1–95 residues) were expressed in HEK293 cells using a pCDNA3.1 vector. To ensure that neither an N-terminal nor a C-terminal translocation signal would be disturbed, no tags were fused to the construct, but an antibody was raised against recombinant MOCS1BΔ1–95. HEK293 cells were harvested after two days of protein expression and fractionated into cytosolic (mitochondria free) and non-cytosolic fraction (containing mitochondria). Comparative western blot analysis of the obtained fractions revealed a strong enrichment of MOCS1B to the non-cytosolic fraction while the majority of MOCS1B∆1–95 was retained in cytosolic fraction confirming the presence of an internal translocation signal at the N-terminus of MOCS1B (Fig. 5B), which was in accordance with the loading controls VDAC (mitochondria) and gephyrin (cytosol). The cellular distribution of both MOCS1B variants was subsequently confirmed by transfecting COS7 cells with plasmids expressing both MOCS1B and MOCS1BΔ1–95 as EGFP fusion proteins. While the MOCS1B-EGFP fusion protein again localized to mitochondria (Fig. 5C), we found that the MOCS1BΔ1–95-EGFP fusion showed a cytosolic distribution of the protein (Fig. 5D), demonstrating indeed the presence of an additional translocation signal in the exon 10 encoded linker region connecting the MOCS1A-and MOCS1B-domains.

**Figure 5.**
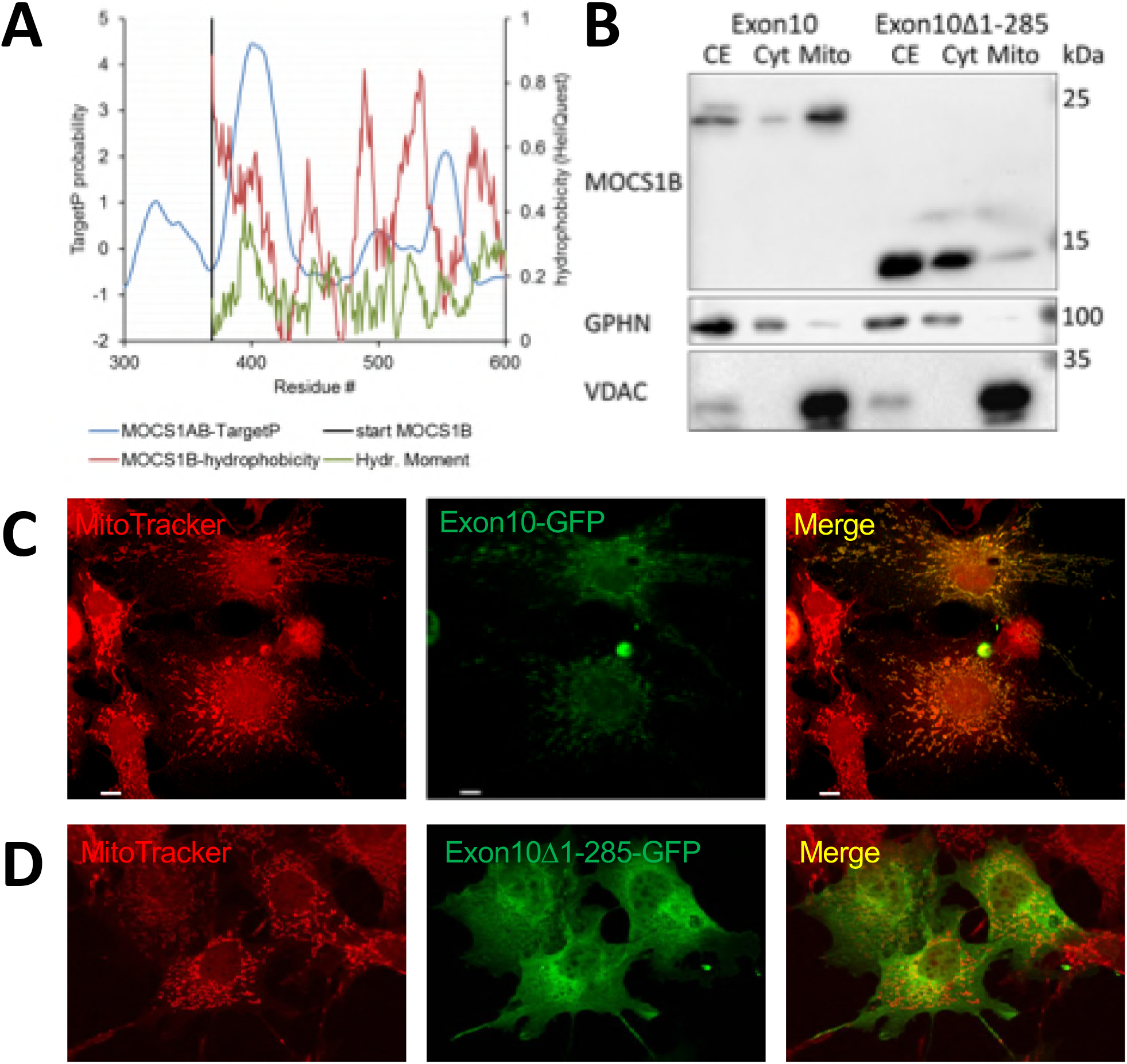
Identification of exon 10 encoded MOCS1B translocation signal. A) TargetP and heliquest in silico analysis of the MOCS1AB amino acid sequence showing the C-terminal residues 300 to 600. B) Cellular fractionation of HEK293 cells transiently expressing the exon 10 encoded either full-length MOCS1B protein or a truncated MOCS1B protein not containing the N-terminal 95 residues. Western blot analysis following 12%SDS-PAGE was performed using anti-MOCS1B, anti-gephyrin and anti-VDAC antibodies. C,D) Transient expression of C) exon 10 and D) exon 10∆1–285 as GFP fusions in COS7 cells. Following 48 h of transient expression, mitochondria were stained with MitoTracker®Red CMXRos and analyzed by confocal laser scanning microscopy. Overlay between the red and green (GFP) channel is shown in the yellow merge panel.

### Sub-mitochondrial localization of MOCS1AB

Considering that MOCS1A proteins are either cytosolic or mitochondrial matrix proteins, we next investigated the sub-mitochondrial localization of MOCS1AB proteins and whether this is influenced by their exon 1 composition. Therefore we first compared the mitochondrial distribution of MOCS1A-ad and MOCS1AB-bcd by expressing both proteins as N-terminal fusions to ratiometric pHluorin (Miesenbock et al., 1998) in HEK293 cells.

First we calibrated the ratiometric pHluorin protein to different pH values. Cells were harvested, fractionated and finally the non-cytosolic fractions were resuspended in buffers of different known pH-values and disrupted resulting in excitation spectra differing in their excitation maxima at 388 nm and 456 nm in a ratiometric manner (Fig. 6A). As a result, the excitation peak at 388 nm was rising with increasing pH, while the signal at 456 nm was decreasing accordingly. The ratio of the excitation bands (388 nm/456 nm) was than related to the respective pH-values and a sigmoidal fit curve was applied (Fig. 6B) allowing the calculation of an unknown pH-value from determined excitation ratios.

**Figure 6.**
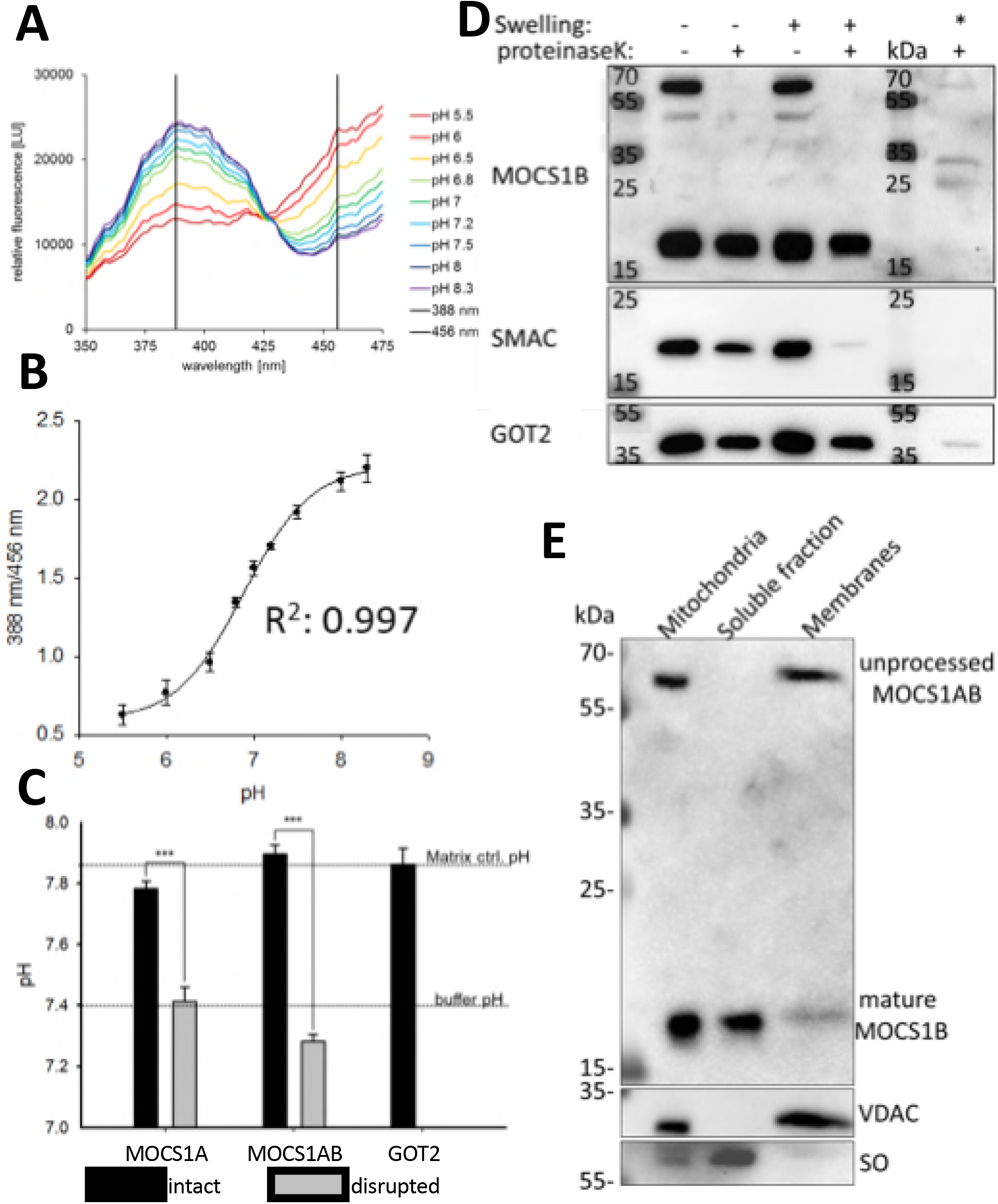
Submitochondrial localization of MOCS1AB proteins. A-C) Excitation-scan measurements of enriched mitochondria from HEK293 cells transiently expressing MOCS1A-ad, MOCS1AB-bcd or GOT2 as pHluorin fusions at 510 nm emission wavelength. A) MOCS1-ad excitation spectra of disrupted mitochondria in buffer of different known pH-values. B) Sigmoidal-fit calibration curve of pHluorin of the excitation maxima ratios (388 nm/456 nm) versus the pH-values. C) Fluorescence based pH-determination of intact mitochondria (black bar) and disrupted mitochondria (grey bar) using the pHluorin calibration curve. D) Partial proteolysis experiment of mitochondria enriched from HEK293 cells overexpressing MOCS1AB-ad without tag. One of each fraction of intact and swollen mitochondria were treated with proteinase K, as well as one additional sample of disrupted mitochondria. Subsequent 12%-SDS-PAGE followed by western blot analysis was performed using anti-MOCS1B, anti-SMAC and anti-GOT2 antibodies. E) Western blot analysis of sub-fractionated HEK293 cells. Mitochondrial membranes and soluble mitochondrial fractions were separated by alkaline extraction and subsequent ultra-centrifugation. Successful separation was demonstrated using anti-sulfite oxidase and anti-VDAC antibodies. Unprocessed MOCS1AB and mature MOCS1B were visualized using anti-MOCS1B antibody.

Next, both MOCS1A-ad-pHluorin and MOCS1AB-bcd-pHluorin were expressed in HEK293 cells, harvested and fractionated. The non-cytosolic fraction was again resuspended in a buffer of known pH (7.4), but was left intact. Measurement of the intact mitochondria yielded pH-values of 7.78±0.02 for MOCS1A-ad and 7.90±0.03 for MOCS1AB-bcd indicating matrix localization for both proteins when compared to the mitochondrial matrix protein GOT2 (7.86±0.05). Following the measurement of the intact mitochondria, samples were disrupted and re-measured resulting in the expected pH-values of 7.41±0.05 and 7.28±0.02 for MOCS1A-ad and MOCS1AB-bcd resembling the pH of the used buffer and differing from the pH of the intact mitochondria significantly (Fig. 6C).

In addition to the ratiometric pHluorin studies, localization of MOCS1A-ad and MOCS1AB-bcd protein to the mitochondrial matrix strongly suggested matrix localization of MOCS1AB-ad as well. To confirm this proposal, MOCS1AB-ad was expressed in HEK293 cells without any tag using the pCDNA3.1 vector. Following enrichment of mitochondria the aforementioned proteinase K treatment was performed. Surprisingly MOCS1AB could not be detected by the MOCS1B antibody in either sample that was exposed to proteinase K, but it was present in both control samples (Fig. 6D), suggesting that MOCS1AB is localized to the outer side of the outer mitochondrial membrane. The experiment was also repeated using EGFP fused MOCS1AB-bcd (Fig. EV4), with a similar result indicating that neither the EGFP nor the pHluorin tag, nor the exon 1 composition alters the translocation of MOCS1AB proteins. This finding was in contrast to the obtained results of the pHluorin measurements.

**Extended View figure 4.**
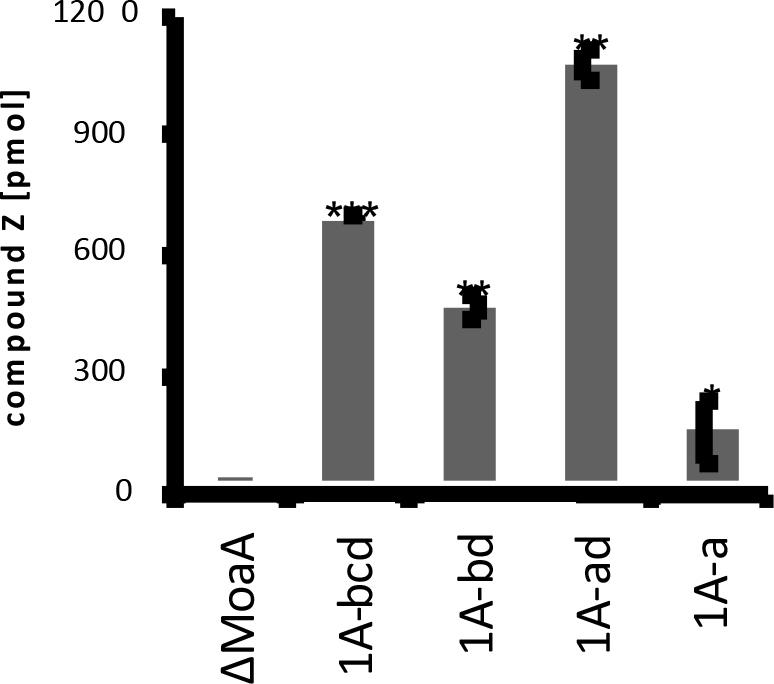
Partial proteolysis experiment of mitochondria enriched from HEK293 cells. HEK293 cells overexpressing MOCS1AB-bcd-MYC (A) or MOCS1AB-ad-GFP (B) fusion proteins were fractionated and partial proteolysis was performed. One of each fraction of intact and swollen mitochondria were treated with proteinase K, as well as one additional sample of disrupted mitochondria. Subsequent 12%-SDS-PAGE followed by western blot analysis was performed using anti-SMAC and anti-GOT2 antibodies as loading controls. Tagged MOCS1AB proteins were detected using anti-MYC (A) or anti-GFP (B) antibodies.

In addition to the bands observed for the full-length MOCS1AB proteins, the MOCS1B antibody also detected a band at approximately 20 kDa in both proteinase K treated samples as well as untreated samples, but not in a control sample of disrupted mitochondria treated with proteinase K (Fig. 6D). A comparable 20 kDa band was also detected when we used C-terminal-tag antibody (Fig. EV4). These findings suggests that a MOCS1B protein lacking the entire N-terminal MOCS1A domain, but not full-length MOCS1AB, was imported into the mitochondrial matrix. To provide further support for the exclusive localization of MOCS1B to the mitochondrial matrix, mitochondria were enriched from HEK293 cells overexpressing MOCS1AB-ad and exposed to an alkaline extraction of the mitochondrial membranes with the aim to separate the inner and outer mitochondrial membranes from the soluble fractions consisting of the intermembrane space and the mitochondrial matrix. Subsequent western blot analysis revealed an exclusive localization of full-length MOCS1AB to mitochondrial membranes while the proteolytically processed MOCS1B protein was mainly found in the soluble fraction, which was in line with the respective loading controls for IMS (sulfite oxidase) and outer membrane (VDAC) (Fig. 6E). Therefore the full-length MOCS1AB protein represents the apo-protein of MOCS1B, which is proteolytically cleaved during procession to the mitochondrial matrix N-terminal to the MOCS1B domain, resulting in a mature soluble MOCS1B protein.

## Discussion

In this study we were able to dissect the impact of alternative splicing of both exon 1 and exon 9 of *MOCS1* on the complex maturation path of MOCS1 proteins leading to the mitochondrial localization of MOCS1A and MOCS1B. The major impact of exon 9 splicing on MOCS1 proteins has been known for years (Gray & Nicholls, 2000), defining the expression of two different types of proteins, MOCS1A or MOCS1AB, respectively. The contribution of exon 1 splicing was not understood, except that exon 1d was found to be essential for catalytic activity as it contains two conserved cysteine residues required for the coordination of the N-terminal [4Fe-4S] cluster. This cluster is crucial for catalytic function of MOCS1A as it is assumed to generate an adenosyl radical in the same way as MoaA in bacteria (Hover & Yokoyama, 2015). This proposal is supported by complementation experiments of MoaA-deficient *E. coli* cells demonstrating a loss of catalytic activity in the MOCS1A-a splice variant (Fig. EV2).

In this study, we were able to show that exon 1a and exon 1b determine the cellular localization of MOCS1 proteins. While exon 1a-encoded residues mediated translocation of MOCS1A to the mitochondrial matrix, MOCS1A proteins with an N-terminal extension encoded by exon 1b remained cytosolic. The function of exon 1c remains unknown, however it should be considered that exons 1b-d are not separated by introns (Gross-Hardt & Reiss, 2002), but rather represent 1 exon which may be spliced partially, similar to exon 9. In either case, we observed that exon 1c is neither essential for catalytic activity (Fig. EV2) nor for translocation processes (Fig. EV1).

Most surprisingly, we found that splicing of exon 9, which is known to produce either active MOCS1A or active MOCS1B protein, overrides the cytosolic translocation of the two discovered cytosolic splice variants, in respect to their N-terminal extension. This is a result of a sequence encoded by exon 10 representing a 95 residue linker region separating the conserved MOCS1A and MOCS1B domains, which is found in all type II and type III splice variants of MOCS1. This linker is facilitating import into the mitochondrial matrix and contains a mitochondrial peptidase cleavage site as witnessed in our translocation studies, proteolysis experiments, and bioinformatically predicted (using the MOCS1B sequence) by Mitoprot (Claros & Vincens, 1996) at position 437 (MOCS1AB splice variant III). The remaining MOCS1B protein (residue 438–620) has a calculated molecular mass of 19.2 kDa, exactly matching the observed size of the cleavage product. Consistently, the recombinantly expressed MOCS1B protein (missing the entire 95 residue linker region) was observed at 18.4 kDa, while the cleavage product was slightly larger due to the deletion of only 70 N-terminal residues (Fig. EV5). A translocation signal and cleavage site prediction using Mitofates (Fukasawa et al., 2015) displayed a mitochondrial processing peptidase (MPP) cleavage site at position 432 (numbering according to splice type III).

**Extended View figure 5.**
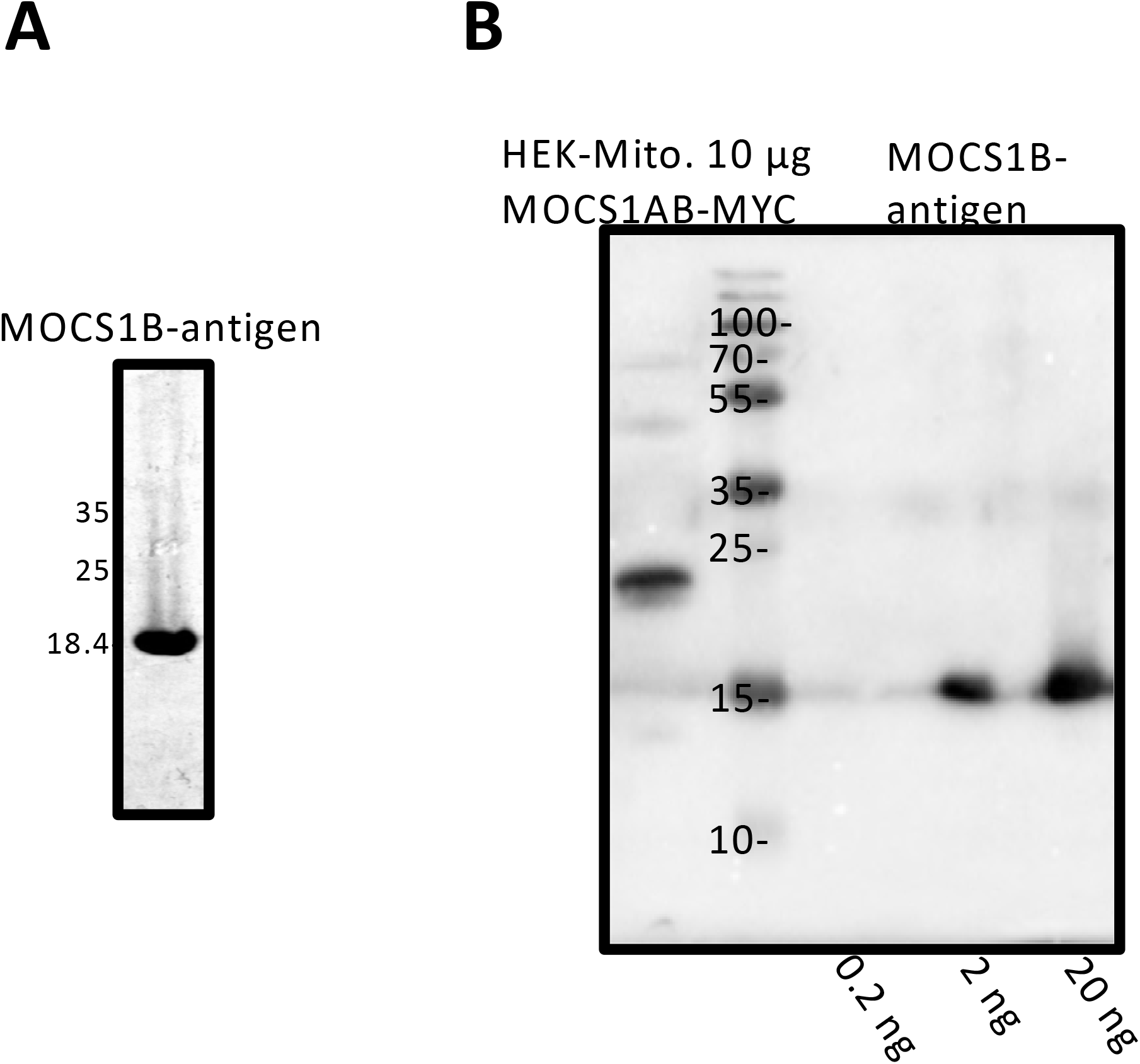
MOCS1B antigen and antibody. A) Coomassie-stained 12% SDS-PAGE of recombinantly expressed, purified His-tagged MOCS1B domain (exon 10 missing 5’ 285 bases) serving as MOCS1B antigen B) WB of 12% SDS-PAGE of 10 µg enriched mitochondria from HEK293-cells overexpressing MOCS1AB and of purified MOCS1B antigen (0.2 ng – 2 ng).

Our findings promote the idea that ancestral *MOCS1A* and *MOCS1B* genes, both encoding their separate localization signals, were fused in vertebrates resulting in an internalization of the MOCS1B translocation signal. As a consequence, we favor the hypothesis of a TOM/TIM23- (Kang et al., 2018) mediated import of MOCS1AB proteins. We furthermore found that exon 1 splicing does not impact MOCS1B procession as observed in partial proteolysis experiments of different MOCS1AB splice variants (Fig. EV4). This is remarkable in light of the competing nature of additional translocation signal present in MOCS1AB-a/ad variants, which in turn suggests a hierarchical import. The fate of the N-terminal MOCS1AB cleavage product consisting of MOCS1A (missing the catalytically essential C-terminal double glycine motif encoded by exon 9) remains unknown.

A very recently reported MoCD type A patient case demonstrated that the expression of the MOCS1AB fusion protein is not strictly required for the biosynthetic activity of MOCS1 proteins. In this patient low amounts of MOCS1B protein (residues 425–620 in splice type III) were able to produce approximately 1% of wildtype Moco levels leading to a mild patient phenotype (Mayr et al., 2018).

In humans, all four genes encoding for Moco-synthetic proteins represent gene fusions (*MOCS1, MOCS2, MOCS3* and *GPHN)*. While *MOCS2* encodes for a unique bicistronic mRNA with overlapping reading frames leading to the orchestrated expression of MOCS2A and MOCS2B proteins by a leaky scanning mechanism resulting in the assembly of the heterotetrameric MPT-synthase complex (Reiss et al., 1999), *MOCS3* and *GPHN* encode for fusion proteins. MOCS3 is responsible for the reactivation of MOCS2A and harbors beside the NifS-like sulfurtransferase domain an additional rhodanese-like domain not present in prokaryotes (Matthies et al., 2004). Gephyrin combines two catalytic activities that are represented by two different Moco-synthetic proteins (MogA and MoeA) (Belaidi & Schwarz, 2013). Orchestrated expression of proteins catalyzing subsequent reaction steps results in the formation of heterooligomers or fusion proteins, which ensures equal expression levels of cooperating proteins (MOCS2) or catalytic domains (MOCS3, gephyrin). MOCS1 represent an example that goes beyond the functional interaction by ensuring orchestrated maturation of MOCS1A and MOCS1B.

In aggregate, biosynthesis of Moco is an evolutionary ancient pathway with each of its reactions steps representing a complex chemical transformation. In particular the reaction mechanism of cPMP formation represents a radical based rearrangement reaction being entirely conserved in all kingdoms of life (Schwarz et al., 2009). Even though the underlying chemistry is highly conserved, the structures of the involved genes differ greatly. Tracking back the evolutionary event of the *MOCS1* gene fusion by aligning MoaA and MoaC in selected species revealed that the gene fusion was not just gained in ophisthokonta, but is a shared feature of the unikonta, given that the amoeba *Dictyostelium discoideum* harbors a fused *MOCS1* orthologue. Therefore the gene fusion event can be traced back to the unikonta/bikonta junction. Besides the above discussed functional benefits of gene fusions in Moco biosynthesis, in the case of *MOCS3* and *GPHN* gene fusion served as evolutionary root for novel functions in higher eukaryotes (Fritschy et al., 2008, Judes et al., 2015).

In conclusion, in this study we were able to demonstrate that MOCS1 proteins target to the mitochondrial matrix. We revealed the effects of alternative splicing on MOCS1 proteins allowing a first comprehensive understanding of the interplay of alternative splicing, cellular translocation and proteolysis mediated maturation of MOCS1 proteins.

## Material and methods

### Materials, plasmids and bacterial strains

Oligonucleotides for PCR and sequencing were purchased from Sigma. Restriction enzymes required were purchased from Fermentas. The T5 RNA polymerase-based bacterial pQE-80L expression vector was used for expression of *MOCS1* splice variants in *E. coli* and purchased from Qiagen. Mammalian expression vectors used for localization studies in COS7 cells and expression in HEK293 cells were pEGPF-N1 and ppHluorin-N1 (derived from pEGFP-N1) and pcDNA3.1myc/His-A, respectively, were from Invitrogen. Bacterial strains *E. coli* wild type MC4100 (*araD139 Δ(argF-lac)U169 rpsL150 relA1 flbB3501 deoC1 ptsF25 rbsR*) and *moaA^-^* and *moaC^-^* mutant strains (*F-thr, leu his pro arg thi ade gal lacY malE xyl ara mtl str Tr λr*) were used for complementation studies.

### Cloning of *MOCS1* splice variants

Constructs were derived from pRitaIX (kindly provided by J. Reiss, Goettingen) (Kugler et al., 2007) while exon 1a was synthesized *in vitro* (Genescript). Based on the published gene sequences (Reiss et al., 1998b, Gross-Hardt & Reiss, 2002, Arenas et al., 2009, Reiss & Hahnewald, 2011) the different splice variants were obtained by two-step PCR reactions fusing the respective exons. The identities of all generated constructs were confirmed by sequencing (GATC-Biotech/Eurofins Genomics).

### Transfection of COS7 and HEK293 cells and transient protein expression

COS7 and human embryonic kidney (HEK293) cells were grown in Dulbecco modified essential medium supplemented with 10% fetal calf serum and L-glutamine. For transfection, cells were seeded into 12-well plates (40,000 cells/well) or 10 cm dishes (4.5 × 10^6^ cells/dish), transfected with Fugene (Promega) or polyethylenimine (PEI, Sigma), respectively, according to manufactures instructions. Transfected HEK293 cells were harvested and prepared for further analysis as described.

### Fluorescence and co-localization analysis

COS7 cells were grown on collagen-coated cover slips, transfected with the respective transgene and cultured for 24 h. Cells were washed with PBS and mitochondria were stained with Mitotracker^®^Red CMXRos according to the manufacturer’s protocol. Following Mitotracker^®^Red staining, cells were washed twice with PBS and fixed with 4 % paraformaldehyde. Coverslips were mounted on microscope slides using Mowiol (Calbiochem) with 1–4-Diazabicyclo(2,2,2)-octan (DABCO, Merck) and dried at 37 °C over night. GFP and Mitotracker^®^Red fluorescence was visualized by confocal laser-scanning microscopy with Tι-Eclipse (Nikon).

### Enrichment and sub-fractionation of mitochondria from transfected HEK293 cells

Mitochondria were enriched from HEK293 cells expressing the respective *MOCS1* splice variants using a modified previously described method (Mattiazzi et al., 2002). All fractions were sonicated for 30 seconds at 10% amplitude using a digital sonifier (Branson, Model 250-D) and centrifuged at 20000 × g for 5 min at 4°C before determination of protein concentration (Roti®-Quant; Roth, Germany) and supplementation with SDS loading buffer to remove protein aggregates. For partial proteolysis experiments enriched mitochondria were separated in 100 µg pellets, which were subsequently resuspended in 100 µl 10 mM HEPES buffer pH 7.6 containing 1 mM CaCl_2_. For the samples of intact mitochondria the buffer additionally contained 220 mM mannitol and 70 mM sucrose. After 5 min incubation on ice the control sample was sonicated for 30 seconds at 10% amplitude and 2 µl of a 5 mg/ml proteinase K (Roth, Germany) solution were added to the respective samples (swollen and swollen + sonicated). After additional 10 min incubation on ice PMSF (final concentration 1 mM) was added and proteins were participated using trichloroacetic acid (final concentration 5%) for 10 min on ice.

Membrane and soluble mitochondrial proteins were fractionated by alkaline extraction. Therefore enriched mitochondria were resuspended in 0.1 M Na_2_CO_3_, pH 11.5. After 30 min incubation on ice, mitochondria were centrifuged at 70000 × g for 1 h at 4 °C. Resulting pellets containing the mitochondrial membranes were washed with H_2_O and resuspended in SDS loading buffer. The supernatants were concentrated using a 3 kDa cutoff protein concentrator and supplemented with SDS loading buffer.

### Western blot analysis

Localization of proteins of fractionated HEK293 cells and sub-fractionated mitochondria was analyzed by Western blot following SDS-PAGE and subsequent semi-dry blot. For the detection of MOCS1 proteins the following antibodies were used in the respective dilutions: anti-MOCS1A (rabbit polyclonal, Abcam, ab176989, 1/200), anti-MOCS1B (rabbit polyclonal, Eurogentec, rabbit SY7796, 1/100), anti-GFP-tag (rabbit polyclonal, Invitrogen, A6455, 1/1000) and anti-MYC-tag (mouse monoclonal, cell supernatant 9E10, 1/5). To ensure successful fractionation and sub-fractionation the following antibodies were used as controls: anti-GOT2 (Sigma, rabbit polyclonal, SAB2100–950, 1/500), anti-gephyrin (mouse monoclonal, cell supernatant 3B11, 1/20), anti-SMAC/Diablo (rabbit polyclonal, Abcam, ab8114, 1/2000), anti-sulfite oxidase (mouse monoclonal, Abcam, ab57852, 1/1000) and anti-VDAC1/Porin (rabbit polyclonal, Abcam, ab15895, 1/1000). All antibodies were diluted in TBST containing 2% dried non-fat milk powder.

### MOCS1B antibody

The MOCS1B domain (encoded by exon 10 Δ1–285) was expressed in *E. coli* BL21 as N-terminally His-tagged protein using a pQE80-L vector. Purification was achieved using Ni-NTA resign. The protein was eluted using a potassium phosphate pH 8.0 buffer containing NaCl and imidazole (250 mM). Subsequently the buffer was exchanged to a 20 mM Tris-HCl buffer pH 7.8 containing 200 mM NaCl using a PD10-column (GE Healthcare) and diluted to 10 mg/ml. Purity of the protein was confirmed by a 12% SDS-PAGE using Coomassie-staining. The antibody was subsequently produced using the acquired protein as antigen (Eurogentech) and tested against different antigen concentrations (0.2–20 ng), as well as recombinantly expressed MOCS1AB from HEK293 cell extract.

### Determination of intra-mitochondrial pH-values

Mitochondria were enriched from HEK293 cells overexpressing the desired proteins as pHluorin fusions. For the determination of the pHluorin pKa-curves pellets of enriched mitochondria were resuspended in water (2 mg/ml) and sonicated for 30 s at 10% amplitude. Subsequently, 100 µl extract were mixed with 100 µl 2 × buffer (50 mM MES or Tris-HCl buffer pH 5.5–8.3, 0.3 M NaCl) and excitation spectra were recorded from 350 nm to 500 nm at an emission wavelength of 510 nm at room temperature using a microplate reader (Tecan InfiniteM200). To measure the pH-values of pHluorin fusions in intact mitochondria, pellets of enriched mitochondria were resuspended in buffer (pH7.4 at 37°C; 1 mg enriched mitochondria/ml) and fluorescence was immediately determined as described above, but at 37°C. Subsequently, each sample was sonicated for 30 s (10% amplitude) and remeasured.

### Determination of cPMP and MPT content

cPMP was detected as Compound Z in crude cell extracts as described (Hanzelmann et al., 2002). In brief, protein extracts (0.5 mg total protein) were oxidized, purified by quarternary aminoethyl column chromatography and HPLC analysis (Agilent Technologies) was performed using a C18-reversed phase column (Waters). Elution from quarternary aminoethyl columns was performed with 50 mM HCl. cPMP concentrations were expressed in picomoles of compound Z per milligram of total protein.

### Statistics

If not stated otherwise, all numerical data were reported as mean ± standard deviation. All measurements were performed at least in triplicates. Statistical analysis (unpaired t-test) was performed with SigmaPlot (Systat Software Inc). Significance levels are indicated in figures as \**p<*0.05, \*\**p<*0.01, \*\*\**p<*0.001.

## Acknowledgments

Technical assistance by Monika Laurien (University of Cologne, Germany) is gratefully acknowledged. We thank Dr. Jochen Reiss (University of Goettingen) for providing the plasmid pRitaIX and Prof. Reinhart Krämer for providing the pHluorin sequence. We thank Prof. Johannes Hermann and Prof. Timo Mühlhaus for the iMTS-L propensity analysis. We gratefully acknowledge funding by the Deutsche Forschungsgemeinschaft within the SPP1927 priority program.

## Author Contributions

S.J.M. was involved in cloning of plasmids and cell culture work. S.J.M conducted all western bolt experiments and pHluorin pH determinations, was involved in study design and wrote the paper. J.R. was involved in cloning of plasmids and cell culture work. J.R. performed all microscopy and complementation experiments and was involved in study design. G.S. was involved in study design and wrote the paper.

